# Metformin Alleviates Arthrofibrosis via Fibroblast Metabolic Reprogramming

**DOI:** 10.1101/2022.07.05.498844

**Authors:** Zhenglin Zhu, Shengqiang Gao, Hui Zhu, Yi Chen, Dandong Wu, Zhiyu Chen, Jing Zou, Xiangdong Wu, Ning Hu, Di Chen, Wei Huang, Hong Chen

## Abstract

**Background:** Emerging studies have suggested an essential role of fibroblast metabolic reprogramming in the pathogenesis of arthrofibrosis. The metabolic modulator metformin appears to be a therapeutic candidate for fibrotic disorders. However, whether metformin could alleviate arthrofibrosis has not been defined. In this study we have determined if treatment with metformin has the beneficial effect on arthrofibrosis and its underlying mechanism.

**Methods:** Articular capsule samples were collected from patients with/without arthrofibrosis to perform gene and protein expression analysis. Arthrofibrosis animal model was established to examine the anti-fibrotic effect of metformin. Cell culture experiments were conducted to determine the mechanism by which metformin inhibits fibroblast activation.

**Results:** We found that glycolysis was upregulated in human fibrotic articular capsules. In an arthrofibrosis animal model, intra-articular injection of metformin mitigated inflammatory reactions, downregulated expression of both fibrotic and glycolytic markers, improved range of motion of the joint, and reduced capsular fibrosis and thickening. At the cellular level, metformin inhibited the activation of fibroblasts and mitigated the abundant influx of glucose into activated fibroblasts. Interestingly, metformin prompted a metabolic shift from oxidative phosphorylation to aerobic glycolysis in activated fibroblasts, resulting in the anti-fibrotic effect of metformin.

**Conclusion:** Metformin decreased glycolysis, causing a metabolic shift toward aerobic glycolysis in activated fibroblasts and has beneficial effect on the treatment of arthrofibrosis.

**The translational potential of this article:** The finding of this study demonstrated the therapeutic effect of metformin on arthrofibrosis and defined novel targets for the treatment of articular fibrotic disorders.

## 1. Introduction

Arthrofibrosis is a common disabling fibrotic joint disorder, manifesting as a painful and long-lasting restriction of range of motion (ROM) [1]. Arthrofibrosis can occur in most joints, but more frequently affects the shoulder, elbow, and knee. Because of the importance of joint function in daily activity, arthrofibrosis is a highly frustrating and debilitating disorder. For example, a minor loss of knee extension of 5° could create difficulties in walking, and a 50% loss in elbow motion leading to 80% loss of function [2]. Unfortunately, satisfactory treatment options for arthrofibrosis are still lacking. Conservative treatments including non-steroidal anti-inflammatory drugs (NSAIDs), intra-articular corticosteroid injections, and physical therapy have only limited effect. Surgical interventions may rapidly improve joint ROM but are associated with specific complications [3]. In addition, surgical trauma can trigger inflammatory reactions, giving rise to even more distinct fibrotic changes [4–6]. Thus, safe and effective alternative therapeutics are needed to treat this debilitating joint disorder.

The pathogenesis of arthrofibrosis has not been fully elucidated. However, the studies of other fibrotic diseases provide important insights and clues. After exposure to various stimuli, quiescent fibroblasts differentiate into highly proliferative myofibroblasts, which produce excessive fibers of α-Smooth muscle actin (α-SMA) and collagen type 1, leading to extracellular matrix (ECM) deposition and tissue contraction [7, 8]. Therefore, profibrogenic transformation of fibroblasts is the key event to the development of fibrosis.

Emerging studies have suggested an essential role of fibroblastic glucose metabolism in fibroblast activation [9, 10]. ^8^F-fluoro-deoxyglucose positron-emission tomography (^18^FDG-PET) studies showed that glucose uptake is increased in the fibrotic capsules of frozen shoulder patients and is highly correlated with the disease severity [11, 12]. Several reports have revealed that transforming growth factor-β (TGF-β), a well-established profibrotic cytokine, increases oxygen consumption and cellular levels of TCA cycle metabolites in fibroblasts, suggesting that oxidative phosphorylation (OXPHOS) is crucial to support the energetic demand of activated fibroblasts [13, 14]. Thus, fibroblastic glucose metabolism may be a promising therapeutic target for arthrofibrosis.

Metabolic modulator metformin has long been used as the first-line treatment for patients with type 2 diabetes mellitus because of high efficacy and safety profile of the drug. Several animal studies demonstrated that metformin could treat and prevent fibrosis in preclinical models of kidney, lung, heart, liver, and ovary, possibly through inducing fibroblast metabolic shift [15–24]. However, whether and how metformin could alleviate arthrofibrosis has not been described.

In the present study, we determined the effect of metformin on arthrofibrosis and its underlying mechanism. We demonstrated that key glycolytic enzymes were upregulated in human fibrotic joint capsules. On the other hand, intra-articular injection of metformin was able to mitigate inflammatory reactions, downregulate expression of both fibrotic and glycolytic markers, improve knee ROM, and reduce capsular fibrosis and thickening in an arthrofibrosis animal model. We also demonstrated that metformin effectively inhibited TGF-β-induced fibroblast activation and this inhibitory effect was mediated by decreased glycolysis and metabolic shift toward aerobic glycolysis in activated fibroblasts.

## 2. Materials and methods

### 2.1. Patient samples

Articular capsule tissue was collected in accordance with the protocol approved by the First Affiliated Hospital of Chongqing Medical University institutional review board (no. 2020-130). Written informed consent was obtained from all participants. The fibrotic capsules were collected from arthrofibrotic patients undergoing arthroscopic capsular release (n=8; 6 females and 2 males; mean age 55 ±4 years). The control tissue was obtained from patients with normal articular ROM (n=8; 6 females and 2 males; mean age 59 ±3 years). The collected specimens were immediately transferred to 0.9% saline solution and then transferred to 4% paraformaldehyde solution after trimming for follow-up morphological examination. The protocol used in the present study was compliant with the ethical guidelines of the Helsinki Declaration in 1975.

### 2.2. Animal studies

All animal studies were approved by our Institutional Animal Care and Use Committee. Eight-week-old male Sprague Dawley rats with a mean weight of 250 g were used in the present study. Rats were housed in the standard cages in a temperature-controlled room with a 12 h light/dark cycle. Twenty rats were divided into four equal groups: control group with only skin incision but no knee immobilization (sham, n=5), immobilized group without intra-articular injection (surgery, n=5), immobilized group injected with an empty vehicle (vehicle, n=5), and immobilized group injected with a metformin-loaded vehicle (metformin, n=5). The vehicle used for intra-articular injections in the present study was poly (lactic-co-glycolic acid) (PLGA) (Sigma-Aldrich, MO, USA). To prepare the metformin-loaded vehicle, PLGA was dissolved in CH_2_Cl_2_, mixed with metformin dissolved by sonication, and then subjected to a double-emulsion procedure. The preparation protocol and controlled release characteristics of this metformin-loaded PLGA have been described and previously validated [25, 26].

Immobilization of the right knee was performed by internal fixation of the femur and tibia, as previously described [27–29]. The surgical procedures were performed under general anesthesia with an intraperitoneal injection of ketamine (100 mg/kg) and xylazine (10 mg/kg). After prepping, the knee joint was hyperextended to 45° to disrupt the posterior capsule. Subsequently, a 3 cm long skin incision was made on the anterior surface of the right knee. Muscle bellies adjacent to the mid-diaphysis of the femur and tibia were separated and retracted. Then, a 1-0 Vicryl suture (Ethicon; Johnson & Johnson) was passed several times around both bones and tied to pull the tibia toward the femur, immobilizing the knee joint at 145° flexed position (Suppl-1). Local anesthetic (0.5% ropivacaine) was applied to the surgical site, and the fascial and skin incisions were closed with 4-0 absorbable sutures (Ethicon; Johnson & Johnson). After resuscitation, the animals were left free to move inside the cage. Surgical incisions were observed daily for signs of infection or inflammation. In addition, abnormal or excessive movement of the immobilized limb, which indicated fixation failure, was also monitored.

Four weeks after the surgery, the knee fixation sutures of immobilized rats were cut and removed under general anesthesia. Subsequently, 100 μl PLGA or metformin-PLGA suspension (equivalent to 10 mg/kg metformin) was injected into the operated knee joint in the vehicle group or metformin group, respectively. Four weeks later, the ROM of the right knee joint was measured; then, all rats were sacrificed, and the right knee joints were harvested.

### 2.3. Cell culture and chemicals

Primary human fibroblasts were extracted from normal capsule tissue which was collected during arthroscopic surgery. Isolation and culture of fibroblasts were performed as previously described [30]. Briefly, the capsule tissue was dissected into 1 mm^3^ piece and digested with 0.1% collagenase I (Sigma-Aldrich, USA) at 37 °C for 2 h. After being filtered and centrifuged, the tissue pellets were cultured in Dulbecco’s modified Eagle’s medium (DMEM, Gibco, Invitrogen, Carlsbad, CA, USA) supplemented with 10% fetal bovine serum (FBS) (Gibco) and 1% penicillin-streptomycin (Gibco). All cells were cultured at 37 °C with 5% CO_2_ and passaged every 2-3 days upon reaching approximately 90% confluence.

Metformin (Selleck Chemical, S1950, TX, USA) was dissolved in phosphate buffered saline (PBS). TGF-β was purchased from Peprotech (100-21&300-02, NJ, USA). Lactate was purchased from Sigma-Aldrich (867-56-1, MO, USA). Unless indicated otherwise, all chemicals were purchased from Sigma-Aldrich or Corning.

### 2.4. Cell viability

Cell viability was determined by the CCK-8 method (Med Chem Express, Shanghai, China) following the manufacturer’s protocol. To assess the toxicity of the dosage, primary fibroblasts were treated with a gradient of metformin concentrations (0-8 mM) for 48 h. The optical density (OD) values were determined at a wavelength of 450 nm. Nonlinear regression analysis was used to calculate the half-maximal inhibitory concentrations (percentage of cell proliferation versus drug concentration).

### 2.5. Small interfering RNA (siRNA) transfection

Small interfering RNA targeting LDHA and negative control (NC) siRNA were chemically synthesized by Tsingke Biotech (Tsingke Biotech, China). Transfection of siRNA at a final concentration of 20 nM was performed using Lipofectamine 3000 (Invitrogen, CA, USA) according to the protocol recommended by the manufacturer.

### 2.6. Reverse transcription-quantitative polymerase chain reaction (qRT-PCR)

Total RNA was extracted from the articular capsule tissues and primary fibroblasts using TRIZOL reagent (Thermo Fisher Scientific, MA, USA) according to the manufacturer’s instructions. Reverse transcription was carried out by using an EvoScript Universal cDNA master reagent kit (Med Chem Express, Shanghai, China). SYBR Green qPCR Master Mix (Med Chem Express, Shanghai, China), cDNA, and the primers were mixed according to the manufacturer’s instructions. β-Actin was used as a reference gene. qRT-PCR was performed in three independent replicates. Sequences of the primers are provided in Suppl-2.

### 2.7. Western blotting

Proteins were extracted from the articular capsule tissues and primary fibroblasts using RIPA buffer containing phosphatase and protease inhibitors. Thirty micrograms of protein extract were subsequently used for western blotting according to a standard protocol. Antibodies against Glucose transporter 1 (GLUT1) (ABclonal, A11727), Hexokinase-2 (HK2) (ABclonal, A0994), Phosphofructokinase (PFKP) (ABclonal, A12160), Pyruvate kinase M-2 (PKM2) (ABclonal, A16700), α-SMA (ABclonal, A1011), Fibronectin-1 (FN1) (ABclonal, A7488), LDHA (ABclonal, A0861), and β-actin (Zen Bio, 200068-8F10) were used. The intensity of the protein bands was analyzed by ImageJ software using β-actin as a reference protein.

### 2.8. Measurement of glucose consumption and lactate production

Glucose and lactate levels were determined as described previously using specific assay kits (Solarbio, BC2235& BC2505, Beijing, China) [31]. A plate reader was used to measure the OD of the reaction solutions (SAMLFTA, Gen5 software; BioTek Instruments, Winooski, VT, USA).

### 2.9. Liquid Chromatograph-Mass Spectrometer (LC-MS)

Mass spectrometry analyses were performed using a high-performance liquid chromatography system (1260 series; Agilent Technologies) and mass spectrometer (Agilent 6460; Agilent Technologies). A 10-cm dish of cultured primary fibroblasts was collected; 1.5 mL of chloroform/methanol (2:1, v/v) was added and the cells were vortexed for 1 min and centrifuged at 3,000 rpm for 10 minutes; 800 μl of organic phase was added to a clean tube and dried under a flow of nitrogen. Mass spectrometry assay was performed after the addition of 200 μl of isopropanol/methanol solution (1:1, v:v), and the supernatant was used for analysis. Data analyses were performed according to the instructions provided by Shanghai Applied Protein Technology.

### 2.10. Immunofluorescence assays of the cells

The cells on the coverslips were incubated with 6% normal goat serum and 4% bovine serum albumin in PBS for 1 h at 37 °C. Then, the cells were incubated with anti-FN1 (ABclonal, A7488) and anti-α-SMA antibodies (ABclonal, A1011) overnight at 4 °C. On the next day, fluorescein isothiocyanate (FITC/CY3)-labeled goat anti-rabbit IgG was added as a secondary antibody and incubated for 1 h at room temperature. The stained cells were washed with 1 × PBS 5 times. Finally, the cells were counterstained with 4’,6-diamidino-2-phenylindole (DAPI) and observed by fluorescence microscopy.

### 2.11. Histological analysis

The samples were fixed in 4% paraformaldehyde (Beyotime, Beijing, China), decalcified in 0.5 M ethylenediaminetetraacetic acid (EDTA), and embedded in paraffin. Serial 5-μm-thick sections were obtained and subjected to histological evaluations. Hematoxylin-eosin (HE) and Masson’s trichrome staining were carried out as previously described [32–34]. Each specimen was scored by three orthopedic surgeons following the double-blind principle according to the Osteoarthritis Research Society International (OARSI) criterion for rats [35].

For immunohistochemistry (IHC) staining, deparaffinized sections were immersed in sodium citrate buffer, heated in a temperature gradient for antigen retrieval, soaked in 3% H_2_O_2_ to remove the endogenous peroxidase activity, and blocked with goat serum. Then, the sections were incubated with antibodies against FN1 (ABclonal, A7488), α-SMA (ABclonal, A1011), GLUT1 (ABclonal, A11727), and LDHA (ABclonal, A0861) at 4 °C overnight. A biotin-labeled secondary antibody (ZSGB-BIO, ZLI-9017) and 3,3’-diaminobenzidine (DAB) were used for detection. The anterior and posterior capsule were evaluated for staining-positive cells.

### 2.12. Sample size determinations

For *in vitro* experiments quantified as fold changes or by quantitative analysis, we calculated a minimum sample size (n ≥3) based on three independent experiments. For *in vivo* experiments performed similar to previous experiments, we estimated a sample size n ≥ 4 for an alpha level of 0.05 and a power level of 80% [36, 37].

### 2.13. Statistical analysis

All datasets were compared using GraphPad Prism (GraphPad Software, La Jolla, CA) version 9.0. The data are shown as the mean ± standard error of mean (SEM). For in *vivo* and *vitro* studies, unpaired Student’s t test (for two groups) or one-way analysis of variance (ANOVA) (for multiple groups) followed by the Tukey-Kramer test was used. *P* < 0.05 was considered statistically significant.

## 3. Results

### 3.1. Glycolysis is upregulated in human fibrotic capsules

Characteristic pathology of arthrofibrosis involves contracture of the thickened and stiffened articular capsule. As shown in H&E sections, the most noticeable changes observed in the fibrotic capsules included the appearance of densely packed collagen fibers, fibroblastic proliferation, and a decrease in synovial fat tissue. Increased vascularity and infiltration of inflammatory cells were also detected in the fibrotic tissue but not in nonfibrotic tissue. Similar to these results, Masson’s trichrome staining showed significant fibrosis in the fibrotic capsules (Fig. 1a). The results of gene and protein expression analysis demonstrated increased expression of fibrotic markers, including α-SMA, FN1, and TGF-β, in the capsules from adhesive capsulitis patients (Fig. 1b-g). These findings were consistent with previous reports about the pathophysiology of arthrofibrosis [38–40].

**Fig.1.**
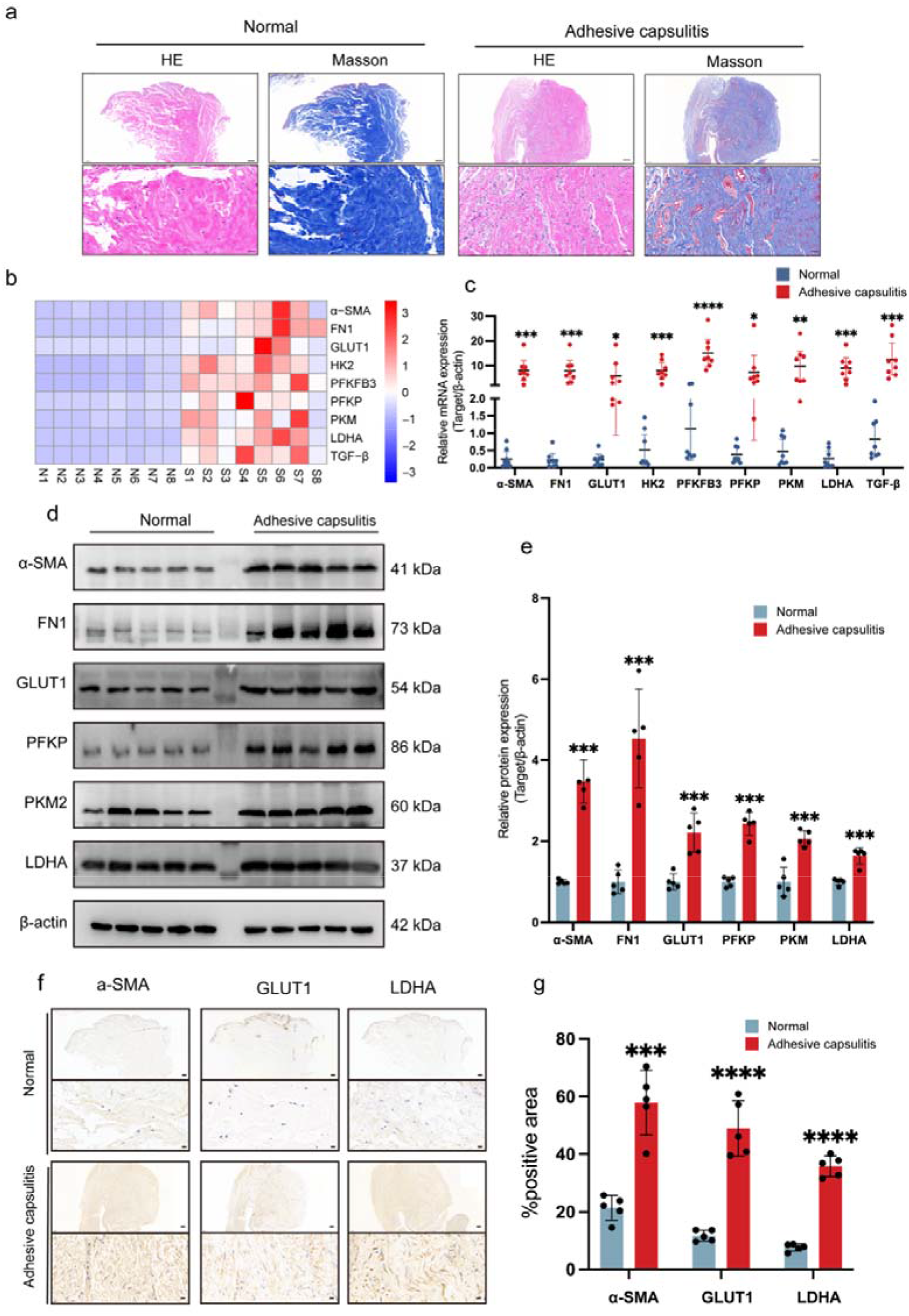
Glycolysis is upregulated in human fibrotic capsules. **a** The capsule tissue samples were obtained from patients with or without shoulder adhesive capsulitis during arthroscopic shoulder surgery and were stained with HE and Masson’s trichrome (n=8, scale bars: upper panel: 200 μm, lower panel: 20 μm). Harvested tissues were homogenized, and mRNA (**b-c**) or protein (**d-e**) levels of α-SMA, FN1, TGF-β, GLUT1, HK2, PFKFB3, PFKP, PKM, and LDHA were quantified by qRT-PCR (n=8) or western blotting (n=5), respectively. The data were normalized to β-actin and expressed as value relative to the level in normal capsular tissue that was designated 1. **f-g** The capsular tissue sections were subjected to IHC staining for α-SMA, GLUT1, and LDHA, and the positive area was quantified (n=5, scale bars: upper panel: 200 μm, lower panel: 10 μm). The data are shown as mean ± SEM. \**P*<0.05, \*\**P*<0.01, \*\*\**P*<0.001, \*\*\*\**P*<0.0001 by unpaired Student’s *t* test.

Next, we measured the expression of the key glycolytic enzymes to assess whether fibrotic capsules are characterized by upregulated glycolysis compared with the nonfibrotic capsules. The data of qRT-PCR analysis indicated that fibrotic capsules exhibited significantly elevated expression of several key glycolytic enzymes, including GLUT1, HK2, PFKFB3, PFKP, PKM, and LDHA (Fig. 1b-c). In agreement with these observations, the data of western blotting showed that the protein expression levels of GLUT1, PFKP, PKM2, and LDHA were significantly upregulated (at least 2-fold) in the fibrotic capsules compared with those in the nonfibrotic capsules (Fig. 1d-e). GLUT1 is responsible for cellular glucose uptake, and abnormal expression of GLUT1 is associated with many disorders [31, 41, 42]. On the other hand, LDHA is critical for the conversion of pyruvate to lactate and dictates the cellular metabolic pathway downstream of glycolysis [43]. Thus, we focused on the expression of GLUT1 and LDHA in the capsule tissues. In agreement with the results of qRT-PCR and western blotting, the data of IHC staining indicated more than two-fold increase in both GLUT1 and LDHA staining positive areas in fibrotic capsules compared with those in normal capsules (Fig. 1f-g). Overall, our data indicated that glycolysis was upregulated in human fibrotic capsules.

### 3.2. Metformin alleviates arthrofibrosis in an animal model

We established a knee arthrofibrosis animal model to explore the anti-fibrotic effect of metformin *in vivo* (Fig. 2a). As shown in Fig. 2b, the operated knee ROM (extension-flexion) was significantly lower in the surgery group than that in the Sham group (−73.0°). No significant differences were detected by comparing the surgery and vehicle groups; however, the ROM of immobilized knees injected with metformin was by 48.6° higher than that of the vehicle-injected knees, indicating that intra-articular injection of metformin significantly improved ROM in the arthrofibrotic knees.

**Fig.2.**
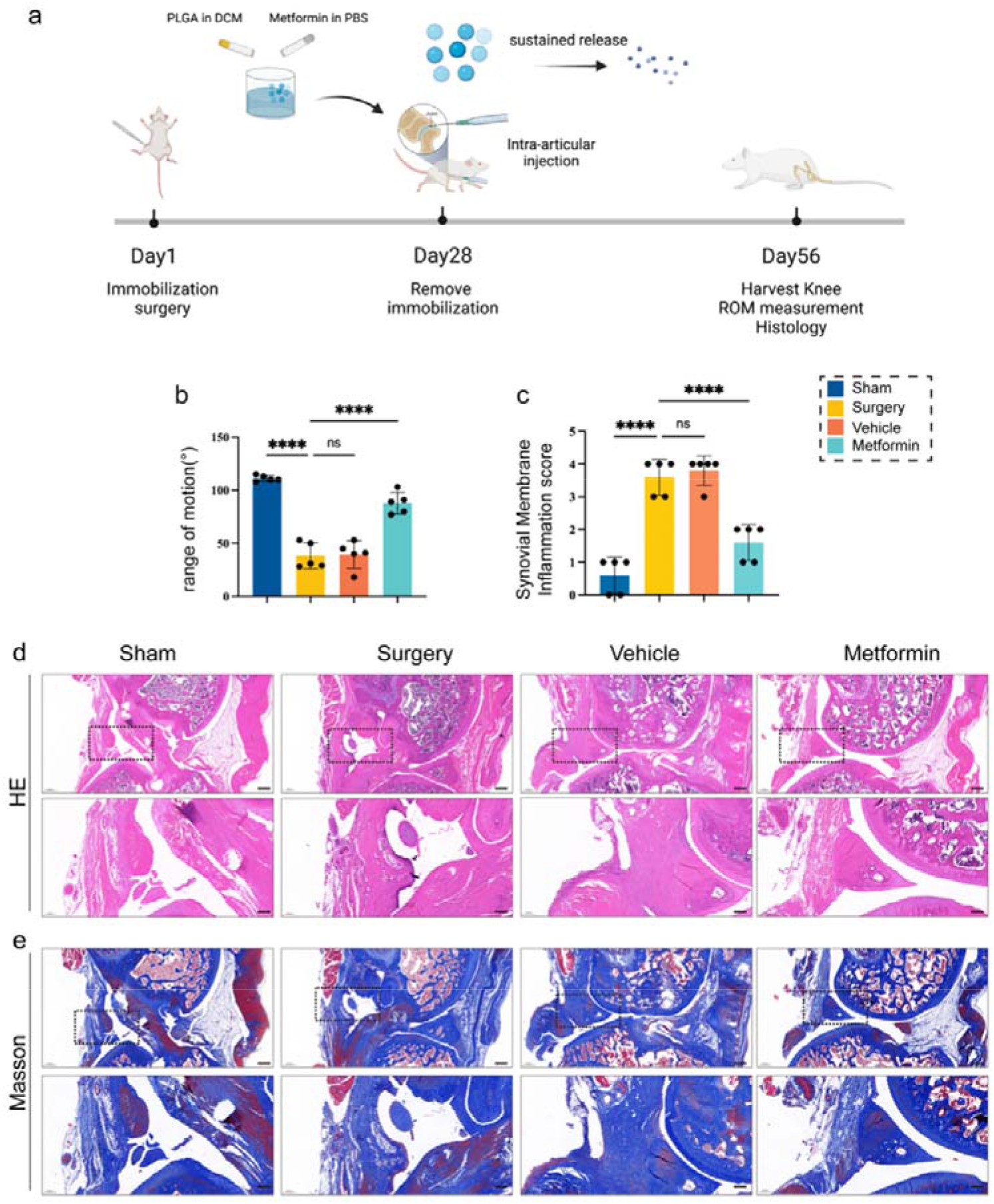
Metformin alleviates arthrofibrosis in animal model. **a** Schematic illustration of the animal study design. Rats were divided into four equal groups: sham, surgery, vehicle, and metformin treatment (n=5). Right knee immobilization surgery was performed on day 1. Four weeks later, the immobilization suture was removed, and metformin or empty vehicle was injected intra-articularly. After another four weeks, the right knee ROM was measured, rats were euthanized, and the knee was harvested for histological examination. **b** ROM measurement at the endpoint of the study (n=5). **c** Synovial membrane inflammation score assessed using HE sections (n=5). **d-e** Immobilized knees were stained with HE and Masson’s trichrome. The data are shown as mean ± SEM. \**P*<0.05, \*\**P*<0.01, \*\*\**P*<0.001, \*\*\*\**P*<0.0001 by one-way ANOVA followed by Tukey-Kramer test. Scale bars: upper panel: 500 μm, lower panel: 200 μm.

We used the synovial membrane histologic inflammation score to evaluate the type, extent, and severity of the inflammation. These properties are based on an increase in the numbers of synovial lining cell layers, proliferation of synovial tissue, and infiltration of inflammatory cells [35]. As shown in Fig. 2c, immobilization induced an obvious intra-articular inflammatory reaction, and metformin treatment produced a significant anti-inflammatory effect. The results of both H&E and Masson staining demonstrated that immobilization surgery led to decrease in synovial layers and synovial fat tissue, prominent capsular thickening, and fibrosis. The vehicle group shared similar histological characteristics with the surgery group; however, the knees of the metformin group were characterized by less severe capsular thickening and fibrosis. In addition, synovial fat tissue was more abundant in the metformin group than that in the surgery and vehicle groups (Fig. 2d-e).

α-SMA is expressed by myofibroblasts and smooth muscle cells of the blood vessels and is a marker of both fibrosis and angiogenesis. On the other hand, FN1 is primarily secreted by fibroblasts and myofibroblasts and has been used as a specific marker of fibrosis [1]. On IHC sections, we found that there were abundant α-SMA-positive and FN1-positive cells at the transition from the capsule to the meniscus (Fig. 3a-b). In the Sham group, α-SMA-positive cells covered 11.1% of the capsule area, and in the surgery group, this number was increased to 48.7%. After metformin treatment, the area of α-SMA-positive cells was significantly decreased to 28.4% (Fig. 3a, e). In the case of FN1-positive cells, the percentage area was 4.3% in the Sham group, 31.1% in the surgery group, and 12.1% in the metformin group (Fig. 3b, f). We also stained the sections with anti-GLUT1 and anti-LDHA antibodies and examined spatial and temporal expression patterns of GLUT1 and LDHA. Similar to the trend observed in the case of α-SMA and FN1, surgery induced a large increase in GLUT1 expression, and metformin injection significantly reduced the percentages of GLUT1-positive cells (Fig. 3c, g). With respect to the LDHA, while surgery group showed large increase in the area of LDHA-positive cells than the Sham group, the area of LDHA-positive cells was even higher in the metformin treatment group (Fig. 3d, h). Overall, these results suggested that metformin mitigated inflammatory reactions, downregulated expression of both fibrotic and glycolytic markers, improved knee ROM and reduced capsular fibrosis and thickening in an arthrofibrosis animal model.

**Fig.3.**
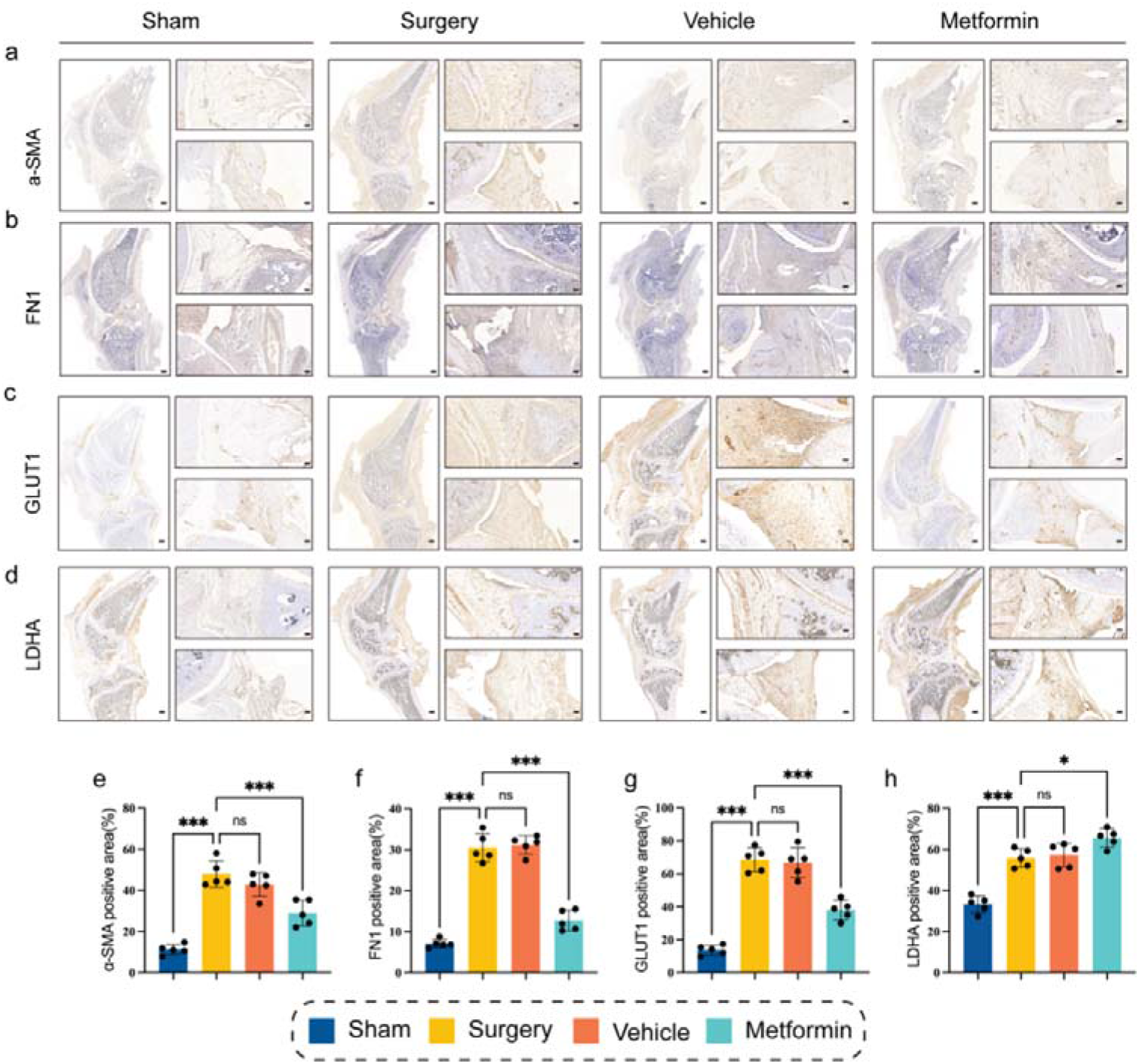
Metformin downregulates both fibrotic and glycolytic markers in an arthrofibrosis animal model. The knee sections of the model animals were subjected to IHC staining for α-SMA, FN1, GLUT1, and LDHA (**a-d**), and the positive area was evaluated (**e-h**). The data are shown as mean ± SEM. \**P*<0.05, \*\**P*<0.01, \*\*\**P*<0.001, \*\*\*\**P*<0.0001 by one-way ANOVA followed by Tukey-Kramer test. Scale bars: left panel: 1000 μm, right panel: 100 μm.

### 3.3. Metformin inhibits the activation of fibroblasts and downregulates fibroblastic glycolysis

The primary human fibroblasts were cultured *in vitro* to determine the influence of metformin on the biological and metabolic properties of fibroblasts. Based on previous reports and the results of the CCK-8 assays performed in the current study, we decided to use 2 mM metformin for the rest of the study [44] (Suppl-3). The data of the immunofluorescence (IF) assay demonstrated that TGF-β, which is the most potent profibrotic cytokine, significantly increased the fluorescence intensity of α-SMA and FN1, and metformin markedly diminished the expression levels of these two fibrotic markers (Fig. 4a-d). These results confirmed that metformin efficiently inhibited TGF-β-induced fibroblast activation.

**Fig.4.**
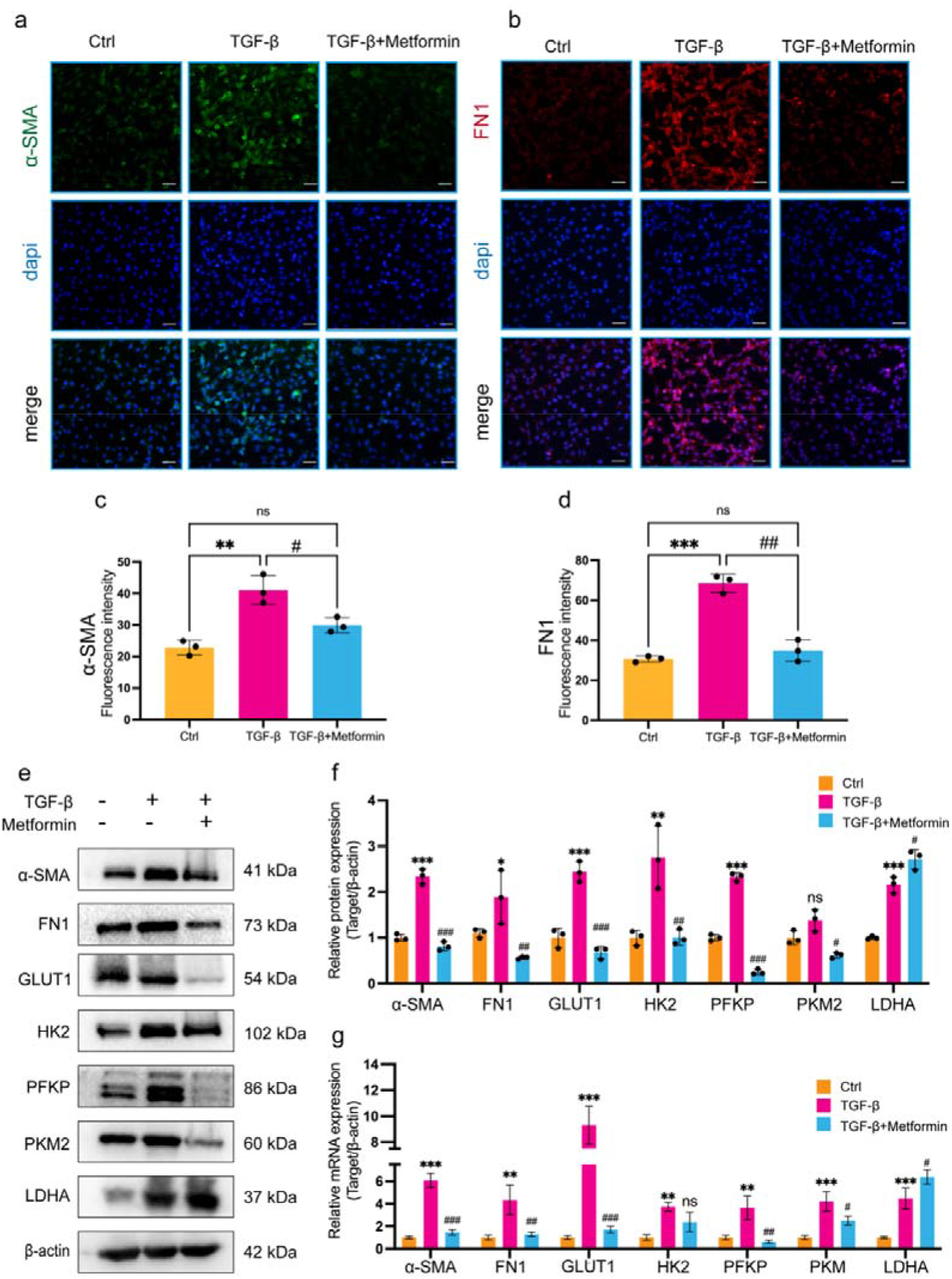
Metformin inhibits fibroblast activation and glycolytic capacity. Primary human fibroblasts were treated with PBS (control), TGF-β (10 ng/ml), or TGF-β (10 ng/ml) + metformin (2 mM) for 48 h. **a-b** Immunofluorescent staining of α-SMA and FN1. **c-d** The results of quantitative analysis showed a significant decrease in α-SMA and FN1 expression in fibroblasts treated with metformin (n=3). **e** The effects of metformin on the protein synthesis of α-SMA, FN1, GLUT1, HK2, PFKP, PKM2, and LDHA. **f** Statistical analysis of protein expression levels shown in (e) (n=3). Protein abundance was normalized to β-actin and expressed as relative value to the level in control cells that was designated 1. **g** mRNA levels of α-SMA, FN1, GLUT1, HK2, PFKP, PKM, and LDHA were quantified by qRT-PCR (n=3). Data was normalized to β-actin and expressed as value relative to the level in control cells that was designated 1. The data are shown as mean ± SEM. *comparison between control and TGF-β treated cells, ^#^comparison between TGF-β and TGF-β + metformin treated cells. \**P*<0.05, \*\**P*<0.01, \*\*\**P*<0.001, \*\*\*\**P*<0.0001 by one-way ANOVA followed by Tukey-Kramer test. Scale bars: 50 μm.

Next, we examined the expression of the key glycolytic enzymes in cultured cells. The mRNA and protein expression levels of the main rate-limiting glycolytic enzymes, including GLUT1, HK2, PFKP, and PKM2, were significantly upregulated by TGF-β, suggesting that fibroblast activation was accompanied by the increased glycolytic activity. These data were consistent with histological findings obtained in human capsular tissue samples. On the other hand, treatment with metformin disrupted the expression of these glycolytic enzymes at both the transcriptional and translational levels (Fig. 4e-g). In contrast to those above-mentioned enzymes, the expression of LDHA was significantly upregulated by metformin. These results, together with data from animal studies, indicate that metformin downregulated fibroblastic glycolysis and acted on metabolic pathways downstream of glycolysis.

### 3.4. Metformin induces metabolic shift in activated fibroblasts

To determine the functional relevance of metformin therapy to glucose metabolism, we analyzed glucose consumption and lactate production in the primary human fibroblasts. TGF-β-treated cells consumed ~80% more glucose than the cells at baseline, and the addition of metformin caused a reduction in glucose consumption of ~25% compared with that in the cells treated with TGF-β alone (Fig. 5a). On the other hand, TGF-β-treated cells produced an increased amount of lactate (~2.5-fold) in the medium, and metformin intervention further promoted lactate production in the context of decreased glucose consumption (Fig. 5b). Consistently, the ratio of lactate production to glucose consumption was the highest in metformin-treated cells (Fig. 5c).

**Fig.5.**
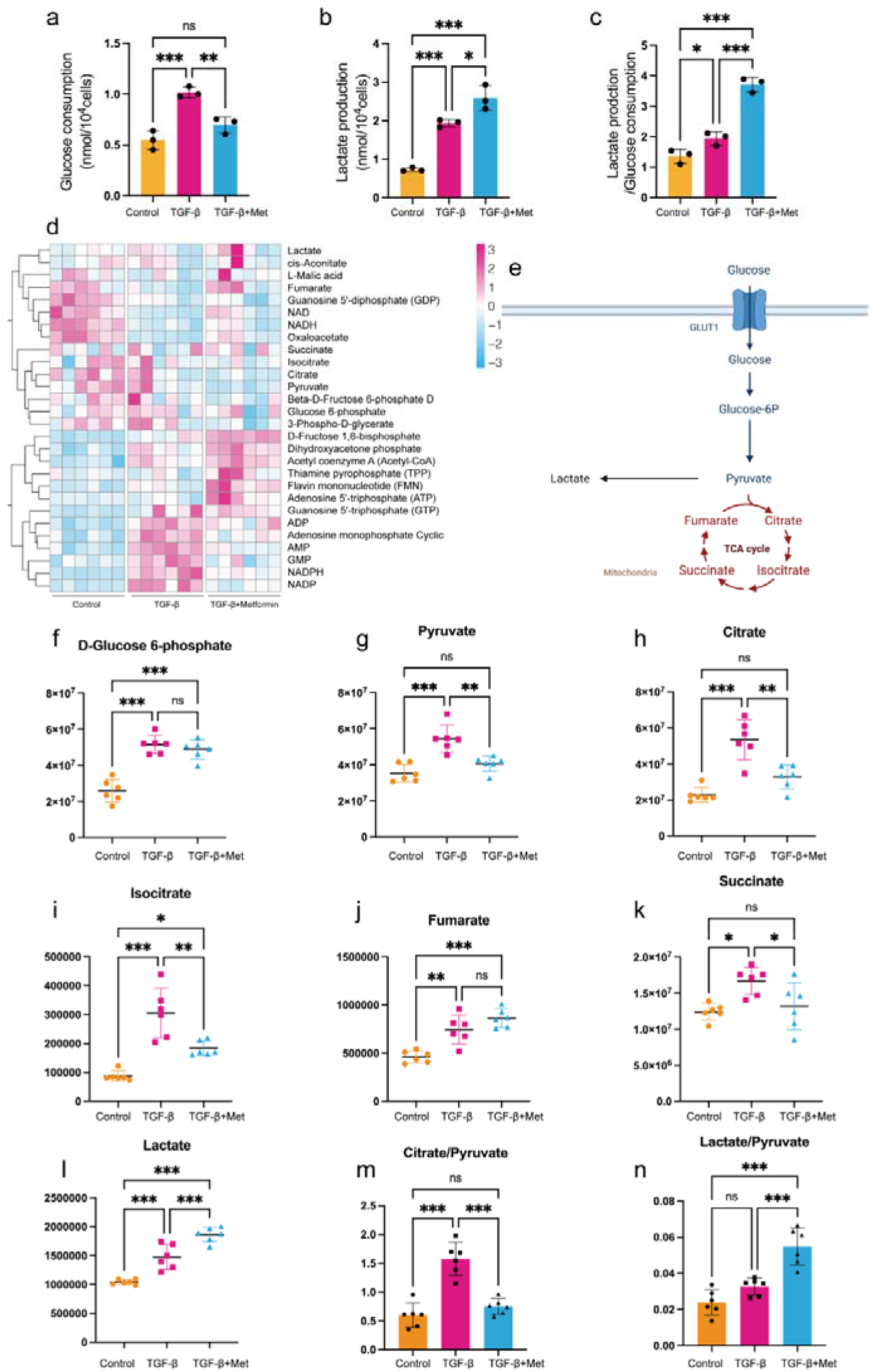
Metformin induces metabolic reprogramming from OXPHOS to aerobic glycolysis in fibroblasts. Primary human fibroblasts were treated with PBS (control), TGF-β (10 ng/ml), or TGF-β (10 ng/ml) + metformin (2 mM) for 48 h. **a-b** Glucose consumption and lactate production by fibroblasts (n=3). **c** The ratio of lactate production to glucose consumption. **d** Heatmap of the main metabolites in control, TGF-β-treated cells, and TGF-β + metformin-treated cells based on LC-MS (n=6). **e** Schematic illustration of the metabolic pathways of aerobic glycolysis and OXPHOS. **f-l** The levels of D-Glucose 6-phosphate, pyruvate, citrate, isocitrate, fumarate, succinate, and lactate in the three groups of cells were compared (n=6). **m-n** Metabolite ratios in differentially treated fibroblasts, including the citrate/pyruvate and lactate/pyruvate ratios (n=6). The data are shown as mean ± SEM. \**P*<0.05, \*\**P*<0.01, \*\*\**P*<0.001, \*\*\*\**P*<0.0001 by one-way ANOVA followed by Tukey-Kramer test.

Accumulating evidence indicates that the effect of metformin is linked to altered metabolic pathways [15, 24, 45–47]. Thus, we systematically assessed the metabolomics profile of fibroblasts using LC-MS analysis after culture with TGF-β with or without metformin (Fig. 5d). Comparison with the cells treated with TGF-β alone indicated that metformin significantly decreased the levels of the main intermediates during the TCA cycle, including citrate, succinate, isocitrate, and fumarate (Fig. 5e-k). In addition, the cellular level of lactate, which is the final product of aerobic glycolysis, was clearly elevated after metformin treatment (Fig. 5l). We also determined the ratios of critical metabolites to better understand the alterations in the metabolic pathways. As shown in Fig. 5m-n, treatment with metformin led to a significant reduction in the citrate/pyruvate ratio and a significant elevation in the lactate/pyruvate ratio, suggesting that metformin stimulated lactate production and inhibited the OXPHOS pathway. In brief, metformin markedly induced a metabolic shift from OXPHOS to aerobic glycolysis in fibroblasts.

### 3.5. Metabolic shift participates in the anti-fibrotic effect of metformin

LDHA is critical for metabolic pathway determination between OXPHOS and aerobic glycolysis. To test the importance of LDHA in the anti-fibrotic effect of metformin, we silenced LDHA in cultured fibroblasts and measured the expression of fibrotic markers. As shown in Fig. 6, treatment with LDHA siRNA significantly increased levels of both the mRNA and protein expression of α-SMA and FN1 in the metformin-treated fibroblasts, indicating that the inhibitory effect of metformin on fibroblast activation was mediated by metabolic shift of fibroblasts.

**Fig.6.**
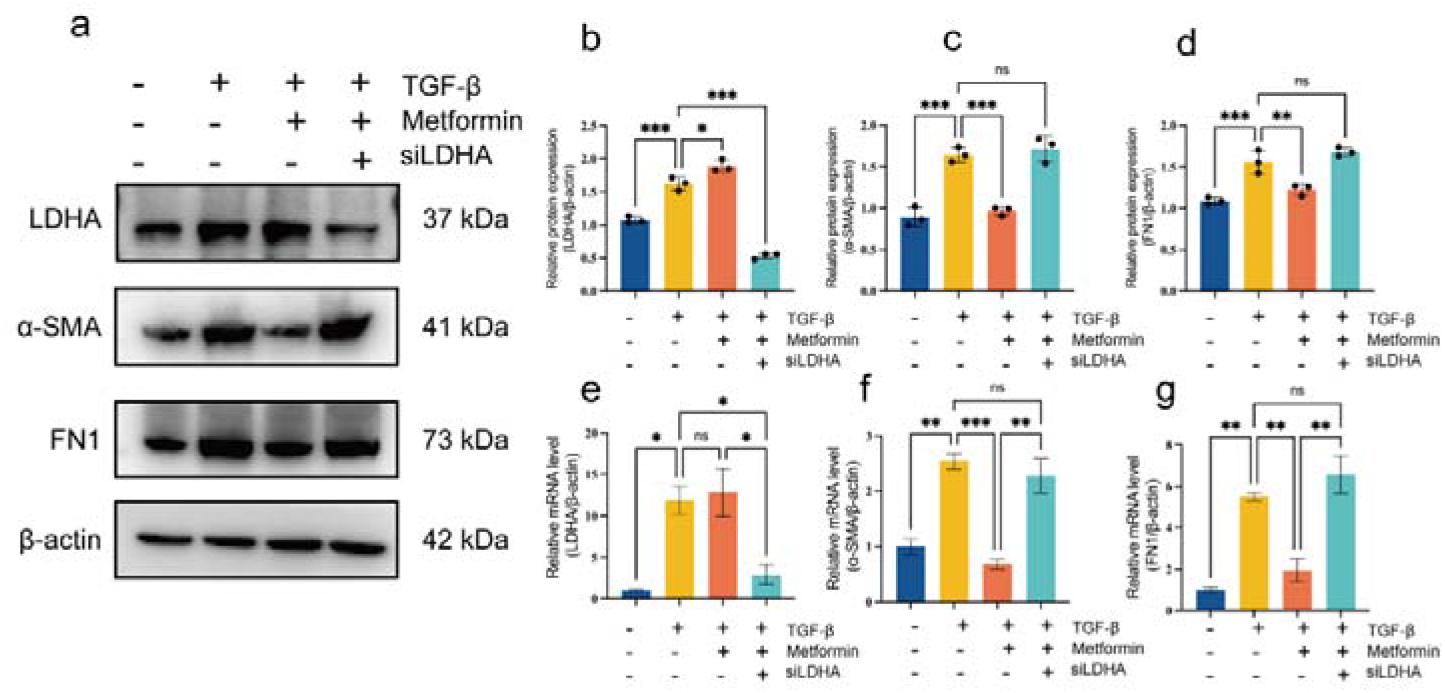
Metformin inhibits fibroblast activation in an LDHA-dependent manner. The primary human fibroblasts were treated with PBS + NC siRNA (control), TGF-β (10 ng/ml) + NC siRNA, TGF-β (10 ng/ml) + metformin (2 mM) + NC siRNA, or TGF-β (10 ng/ml) + metformin (2 mM) + siLDHA for 48 h. **a** Western blot analyses of the total cell lysates of fibroblasts in different groups. **b-e** Statistical analysis of protein expression levels shown in (a) (n=3). Protein abundance was normalized to β-actin and expressed as relative value to the level in control cells that was designated 1. **f-g** mRNA levels of LDHA, α-SMA, and FN1 were quantified by qRT-PCR (n=3). The data were normalized to β-actin and expressed as value relative to the level in control cells that was designated 1. The data are shown as mean ± SEM. \**P*<0.05, \*\**P*<0.01, \*\*\**P*<0.001, \*\*\*\**P*<0.0001 by one-way ANOVA followed by Tukey-Kramer test.

## 4. Discussion

The present study demonstrates the beneficial effect of metformin on arthrofibrosis and its underlying mechanism. Using both the tissue samples and cell culture approaches we demonstrated that the fibrotic phenotype is accompanied by upregulation of glycolytic capacity of fibroblasts. Intra-articular injection of metformin alleviated joint contracture and ECM deposition, mitigated inflammatory reactions, and downregulated both fibrotic and glycolytic markers in a preclinical animal model of arthrofibrosis. At the cellular level, metformin effectively inhibited TGF-β-induced fibroblast activation by decreasing glycolysis and inducing a metabolic shift toward aerobic glycolysis in activated fibroblasts.

The pathogenesis and treatment of tissue fibrosis remain largely unexplored. Emerging studies revealed that dysregulated metabolic changes are involved in the pathogenesis of many fibrotic disorders. Abnormally enhanced OXPHOS activity is one of the major distinguishing phenotypes between activated and quiescent fibroblasts [13, 14]. Cho et al. demonstrated that GLUT1-dependent glycolysis promotes exacerbation of lung fibrogenesis [48]. Upregulation of glycolysis and mitochondrial metabolism is reported to be indispensable to support the high energy demand of liver fibrosis [49]. However, reports regarding the role of metabolic changes in the development of arthrofibrosis are lacking. Our study is among the first to confirm that key glycolytic enzymes are upregulated in both arthrofibrotic patients and animal models. In addition, we have proved that metformin, by inducing metabolic reprogramming in fibroblasts, alleviates the symptoms of established arthrofibrosis in animal models. These results provide evidence for future metabolism-based therapeutics for arthrofibrosis.

Several reports have shown that metformin can prevent and even reverse fibrotic processes in multiple organs, including the kidney, lung, and liver [16–23]. A recent study demonstrated the positive effect of metformin in suppressing joint capsular fibrosis in a mouse model of knee contracture [50]. Previously, the anti-fibrotic action of metformin was reported to be primarily mediated via AMP-activated protein kinase (AMPK) activation and the downstream signaling cascades in fibroblasts, leading to upregulated mitochondrial biogenesis and decreased lactate production [15, 17]. However, a recent metabolomic study has shown that metformin decreases the levels of almost all intermediates of the TCA cycle and increases the amount of lactate in an AMPK-independent manner in fibroblasts [24]. Metformin has also been shown to inhibit Complex I of the respiratory chain, induce lactate production and suppress ATP generation, leading to a metabolic shift from OXPHOS to aerobic glycolysis [46, 47]. The results of the current study are consistent with these findings to some extent, revealing that the metabolic shift toward aerobic glycolysis in fibroblasts played an important role in the anti-fibrotic effect of metformin. The mechanism accounting for these contradictory phenomena remains elusive, implying that the effect of metformin is pleiotropic and highly context dependent [47].

Some limitations exist in the current study. First, the fibrotic shoulder capsule tissues were used to determine the expression of fibrotic and glycolytic markers in arthrofibrotic patients, while a rat model for knee arthrofibrosis was employed for further analysis. Previous clinical and animal studies suggested that fibrotic disorders in different joints shared similar histopathologic and molecular characteristics[2]. Meanwhile, the shoulder capsule samples are more readily available than those from other joints in the clinical setting, while the knee contracture animal model is a well-established and repeatedly validated preclinical model of arthrofibrosis [27–29]. Second, the specific mechanism through which metformin induced metabolic reprogramming in fibroblasts were not determined, which warranted further investigation.

In conclusion, metformin is a potential treatment alternative for arthrofibrosis, the effect of which involves decreased glycolysis and metabolic shift toward aerobic glycolysis in activated fibroblasts (Fig. 7).

**Fig.7.**
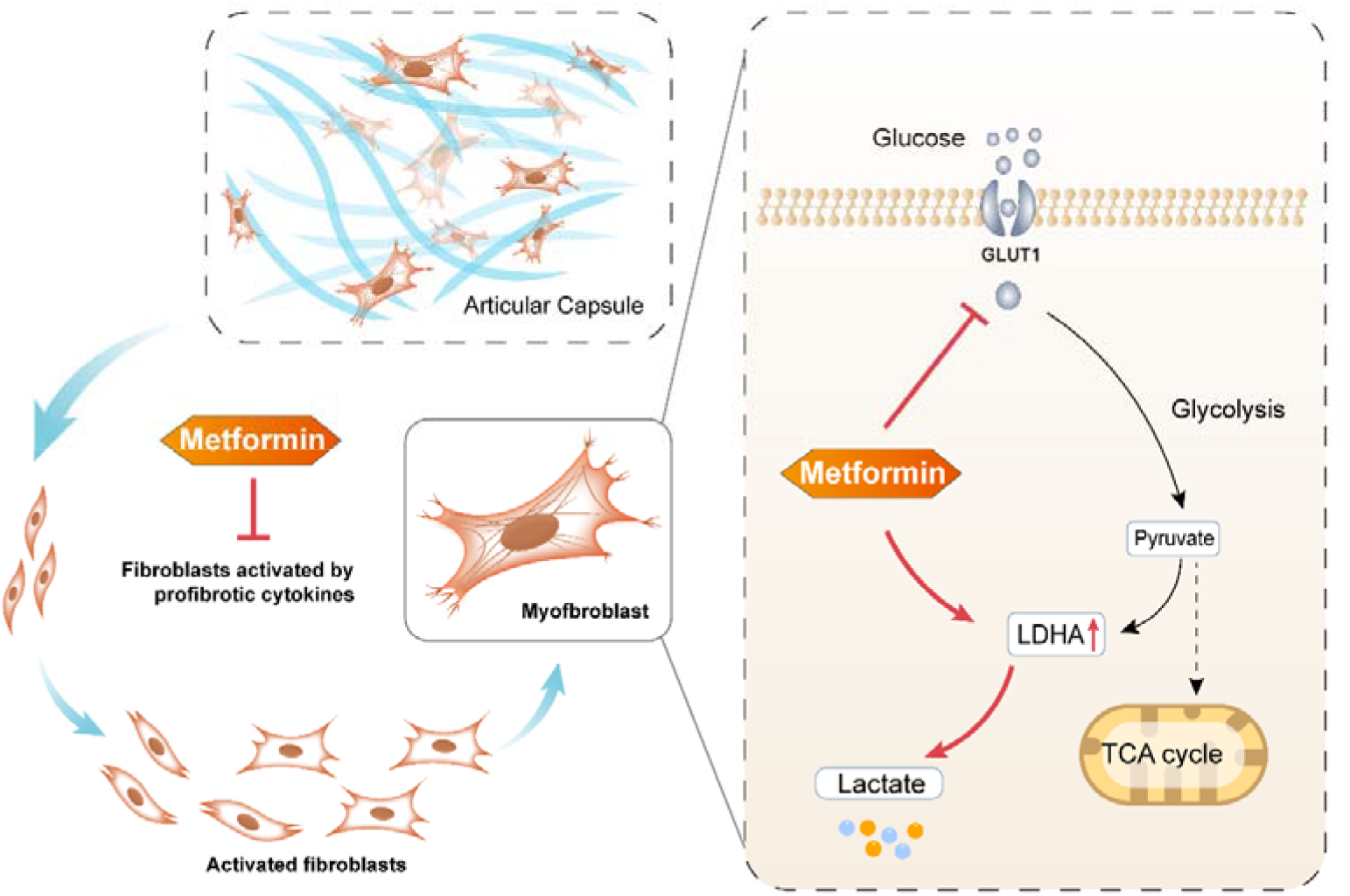
Schematic diagram illustrating the mechanism by which metformin alleviates arthrofibrosis. Metformin effectively inhibits TGF-β-induced fibroblast activation, and this inhibitory effect is mediated by decreased glycolysis and metabolic shift toward aerobic glycolysis in activated fibroblasts.

## Supporting information

Pictures demonstrating the establishment of arthrofibrosis animal model and intra-articular injection.

Primer sequences

CCK-8 assay results to test the most suitable concentration of metformin in in vitro study.

## Descriptive caption for supplementary material

1. Pictures demonstrating the establishment of arthrofibrosis animal model and intra-articular injection.
2. Primer sequences for qRT-PCR.
3. CCK-8 assay results to test the most suitable concentration of metformin in *in vitro* study.

## Acknowledgements

This work was supported by the Chongqing Natural Science Foundation (Grant No. cstc2019jcyj-msxmX0831); the Science and Technology Research Program of Chongqing Municipal Education Commission (Grant No. KJQN202000430); and the Chongqing Medical Scientific Research Project (Grant No. 2021MSXM032).

## Author contributions

**Zhenglin Zhu:** Formal analysis, Investigation, Methodology, Visualization, Writing-original draft. **Shengqiang Gao:** Formal analysis, Investigation, Methodology, Visualization. **Hui Zhu:** Resources, Investigation. **Yi Chen:** Investigation. **Dandong Wu:** Resources, Methodology, Validation. **Zhiyu Chen:** Resources. **Yanran Huang:** Investigation. **Xiangdong Wu:** Methodology, Investigation. **Ning Hu:** Methodology, Validation. **Di Chen:** Writing-review & editing. **Wei Huang:** Conceptualization, Supervision, Writing-review & editing. **Hong Chen:** Conceptualization, Funding acquisition, Writing-review & editing. All authors approved the final version.

