## Supplementary figures and images for "Metformin Alleviates Arthrofibrosis via Fibroblast Metabolic Reprogramming"

### CCK-8 assay results to test the most suitable concentration of metformin in in vitro study.

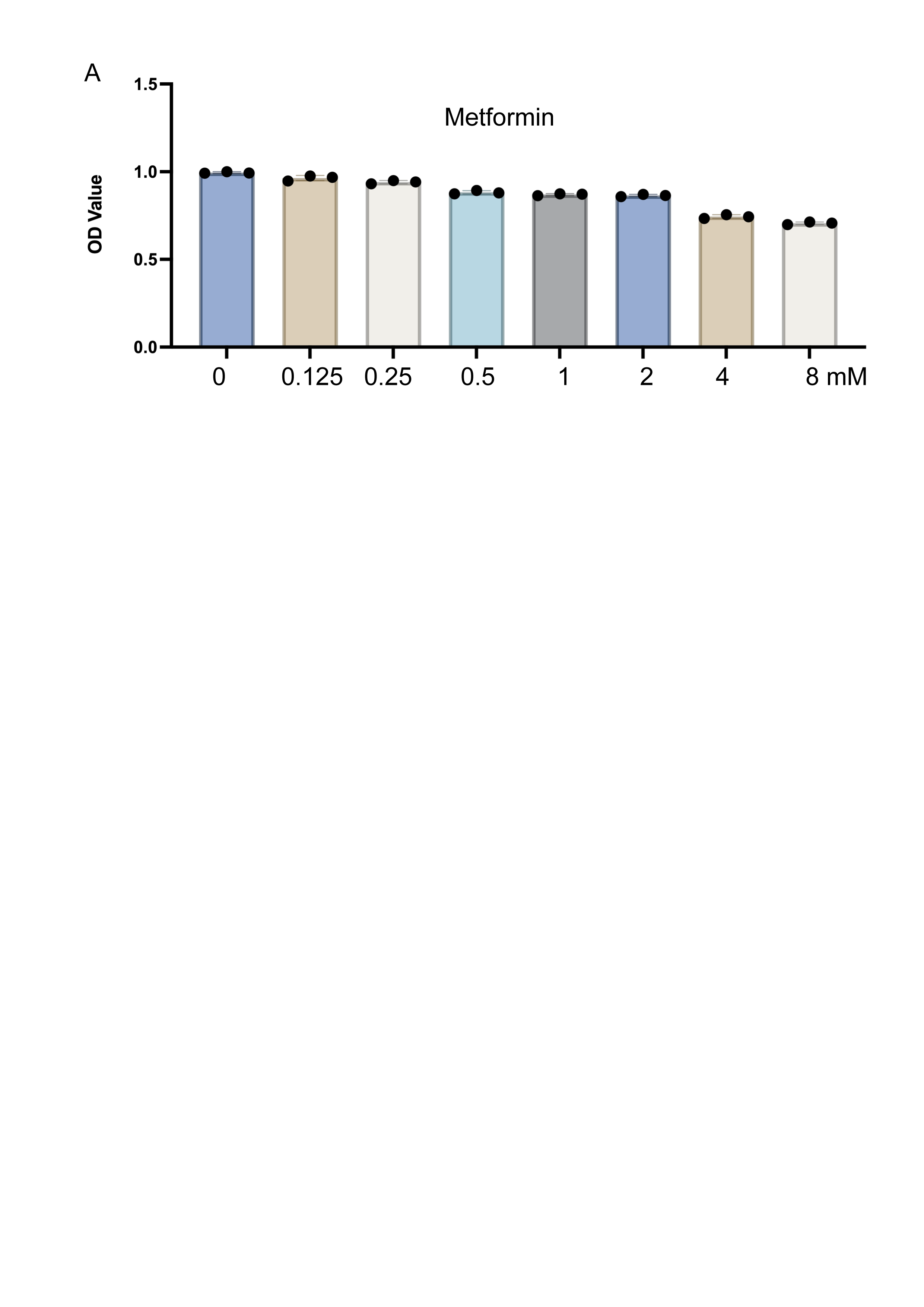

### Pictures demonstrating the establishment of arthrofibrosis animal model and intra-articular injection.

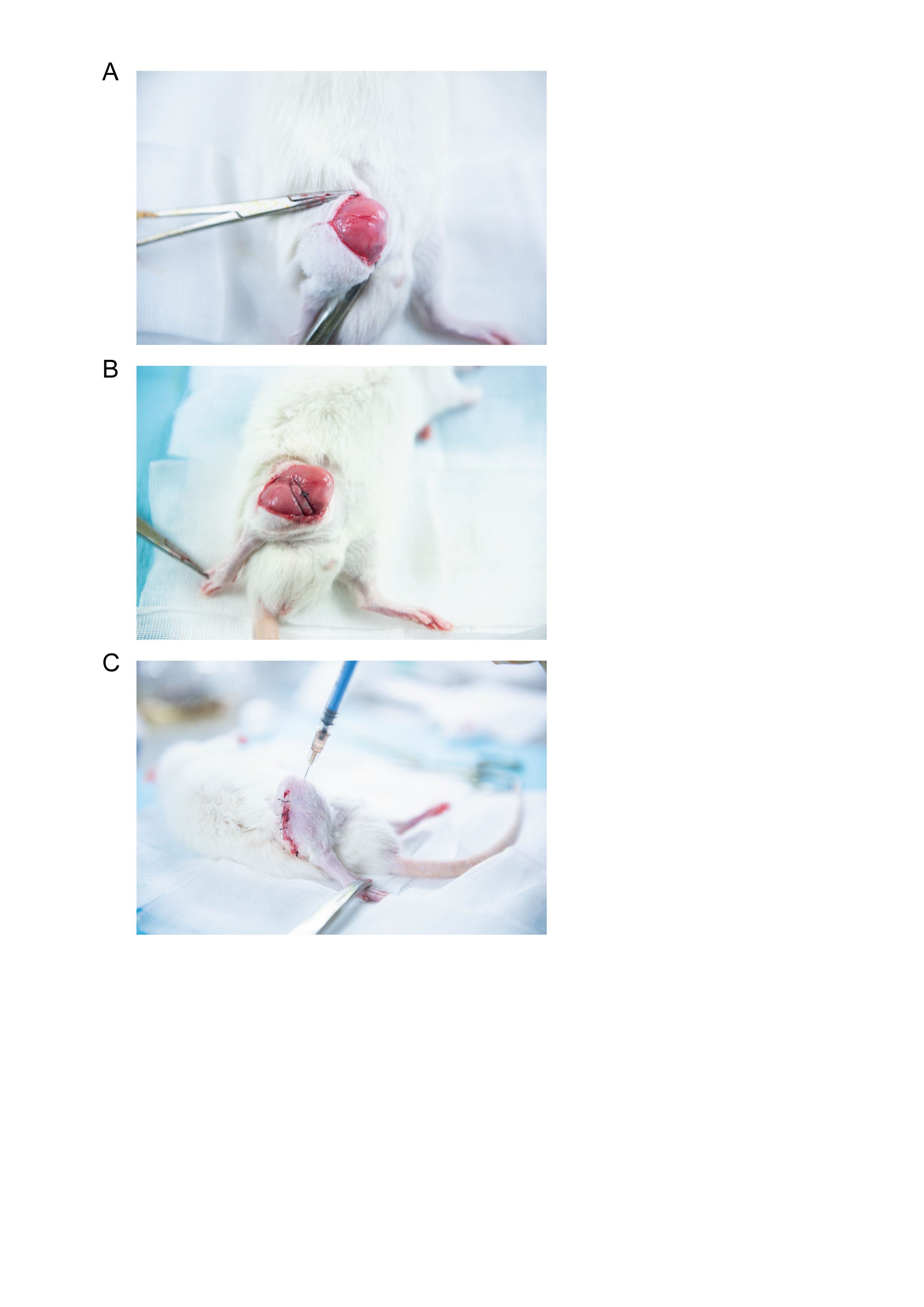
